# Host-virus chimeric events in SARS-CoV2 infected cells are infrequent and artifactual

**DOI:** 10.1101/2021.02.17.431704

**Authors:** Bingyu Yan, Srishti Chakravorty, Carmen Mirabelli, Luopin Wang, Jorge L. Trujillo-Ochoa, Daniel Chauss, Dhaneshwar Kumar, Michail S. Lionakis, Matthew R Olson, Christiane E. Wobus, Behdad Afzali, Majid Kazemian

## Abstract

Pathogenic mechanisms underlying severe SARS-CoV2 infection remain largely unelucidated. High throughput sequencing technologies that capture genome and transcriptome information are key approaches to gain detailed mechanistic insights from infected cells. These techniques readily detect both pathogen and host-derived sequences, providing a means of studying host-pathogen interactions. Recent studies have reported the presence of host-virus chimeric (HVC) RNA in RNA-seq data from SARS-CoV2 infected cells and interpreted these findings as evidence of viral integration in the human genome as a potential pathogenic mechanism. Since SARS-CoV2 is a positive sense RNA virus that replicates in the cytoplasm it does not have a nuclear phase in its life cycle, it is biologically unlikely to be in a location where splicing events could result in genome integration. Here, we investigated the biological authenticity of HVC events. In contrast to true biological events such as mRNA splicing and genome rearrangement events, which generate reproducible chimeric sequencing fragments across different biological isolates, we found that HVC events across >100 RNA-seq libraries from patients with COVID-19 and infected cell lines, were highly irreproducible. RNA-seq library preparation is inherently error-prone due to random template switching during reverse transcription of RNA to cDNA. By counting chimeric events observed when constructing an RNA-seq library from human RNA and spike-in RNA from an unrelated species, such as fruit-fly, we estimated that ~1% of RNA-seq reads are artifactually chimeric. In SARS-CoV2 RNA-seq we found that the frequency of HVC events was, in fact, not greater than this background “noise”. Finally, we developed a novel experimental approach to enrich SARS-CoV2 sequences from bulk RNA of infected cells. This method enriched viral sequences but did not enrich for HVC events, suggesting that the majority of HVC events are, in all likelihood, artifacts of library construction. In conclusion, our findings indicate that HVC events observed in RNA-sequencing libraries from SARS-CoV2 infected cells are extremely rare and are likely artifacts arising from either random template switching of reverse-transcriptase and/or sequence alignment errors. Therefore, the observed HVC events do not support SARS-CoV2 fusion to cellular genes and/or integration into human genomes.

## Introduction

Advances in, and availability of, high throughput sequencing technologies have enabled the accumulation of detailed molecular level information from cells, including genome variations, gene transcription, and gene regulation. These technologies are extremely sensitive at capturing nucleic acid sequences regardless of their origin. As such, the data from these techniques contain not only sequences encoded by the cell itself, but also sequences encoded by infecting pathogens, and/or common contaminating agents (e.g. vectors, plasmids, etc)^1,2^. In virus-infected cells, captured sequences derived from the host or virus represent a powerful tool to study the mechanisms underlying host-pathogen interactions. We have previously deployed these methods to gain mechanistic insights into the pathophysiology of oncogenic viruses, such as Epstein-Barr virus (EBV), hepatitis B virus (HBV) and human papilloma virus (HPV)^1,3–5^.

RNA-sequencing data from virally infected cells contain reads that map perfectly to either the host genome or the viral genome. However, a significant portion of host sequencing reads can also be aligned to discontiguous sections of the genome and often represent canonical forward splicing or back splicing events generated from mRNAs and circular RNAs, respectively. In cells infected with DNA viruses that integrate into the host genome (e.g. HPV or HBV), a chimeric read that is partly mapped to the host genome and partly mapped to the virus genome, is a signature of transcribed segments of the host genome containing integrated viral DNA ^6–8^. Similarly, in virus-induced cancer cells, chimeric reads that are partly mapped to one gene and partly mapped to another gene are the markers of genomic rearrangement and/or gene fusion ^9,10^. Thus, chimeric reads can represent real biological events.

The pathogenic mechanisms underlying severe acute respiratory syndrome coronavirus 2 (SARS-CoV2), the virus responsible for pandemic coronavirus disease of 2019 (COVID-19), are under investigation^11–14^ but still not fully understood. In particular, relatively little is known about the processes following viral infection and why some individuals develop little to no symptoms, while others develop life-threatening or persistent (“long”) COVID-19. Recent studies have identified host-virus chimeric (HVC) reads in RNA-sequencing data from SARS-CoV2 infected cells and samples from COVID-19 patients ^15,16^. Both studies have suggested that HVC events support potential “human genome invasion” and “integration” by SARS-CoV2. This suggestion has fueled concerns about the long-term effects of current vaccines that incorporate elements of the viral genome ^17^. SARS-CoV2 is a positive-sense single-stranded RNA virus that does not encode a reverse transcriptase and does not include a nuclear phase in its life cycle, so some doubts have rightfully been expressed regarding the authenticity of HVCs and the role played by endogenous retrotransposons in this phenomenon. Thus, it is important to independently authenticate these HVC events.

Here we investigated the presence of HVC events in a large number of currently available RNA-sequencing samples from SARS-CoV2 infected cells and patients with COVID-19. Consistent with previous studies, we found that 0.01-1% of viral mapped reads could be characterized as HVC reads. In contrast to reads originating from known or novel splicing junctions, HVC events were not reproduced between different libraries, suggesting that they are either stochastic or artifactual. By counting chimeric events observed in unrelated human RNA-seq libraries prepared in the same manner but containing spike-in of fruit-fly RNA, we estimated that errors in reverse transcription result in ~1% of RNA-seq reads being artifactually chimeric, approximately the same frequency as observed HVCs from SARS-CoV2 infected cells. Finally, we developed a novel experimental approach to enrich for viral sequences from infected cells during RNA-seq library preparation. Despite achieving >30-fold enrichment of viral sequences using our method, HVC events were not enriched, indicating that the low frequency of HVCs observed are likely introduced during reverse transcription steps of RNA-seq library preparation. In summary, we conclude that current data do not support the authenticity of HVC events in SARS-CoV2 infected samples.

## Results

### HVC events are detected in RNA-seq from SARS-CoV2 infected cells but infrequently in samples from patients with COVID-19

RNA-sequencing reads that partly align to the host genome and partly align to the viral genome are the signature of HVC events in RNA-sequencing datasets. Recent reports^15,16^ suggesting the presence of HVC events in SARS-CoV2-infected cells have been interpreted as supporting viral integration into the human genome as a mechanism of viral persistence. To gain insights into the authenticity of these events, we re-analyzed the 3 available RNA-seq datasets from patients with COVID-19 (n=57 samples) and *in vitro* SARS-CoV2 infected cells (n=64 samples). We categorized sequencing reads to those that perfectly aligned to the human genome (build hg38) in a contiguous or discontiguous manner (i.e. reads originating from one exon or reads spanning exon-exon junctions), those that perfectly aligned to the SARS-CoV2 viral genome, and those that partly aligned to both host and viral genomes (potentially representing HVC events) (**Fig. 1A**). Viral mapped reads were detected across several cell lines infected with SARS-CoV2 (**Fig. 1B**). SARS-CoV2 infected Calu-3 and A549-ACE2 cells had the highest percentages (~20-70%) of viral reads, while other cells, including A549 cells and samples from lung autopsies of patients with COVID-19, had dramatically lower representation of viral reads (**Fig. 1B**). The frequency of viral reads in cells infected *in vitro* with other viruses, including influenza A (IAV), middle east respiratory syndrome (MERS) and respiratory syncytial viruses (RSV), were similar (**Fig. S1A**).

**Fig. 1.**
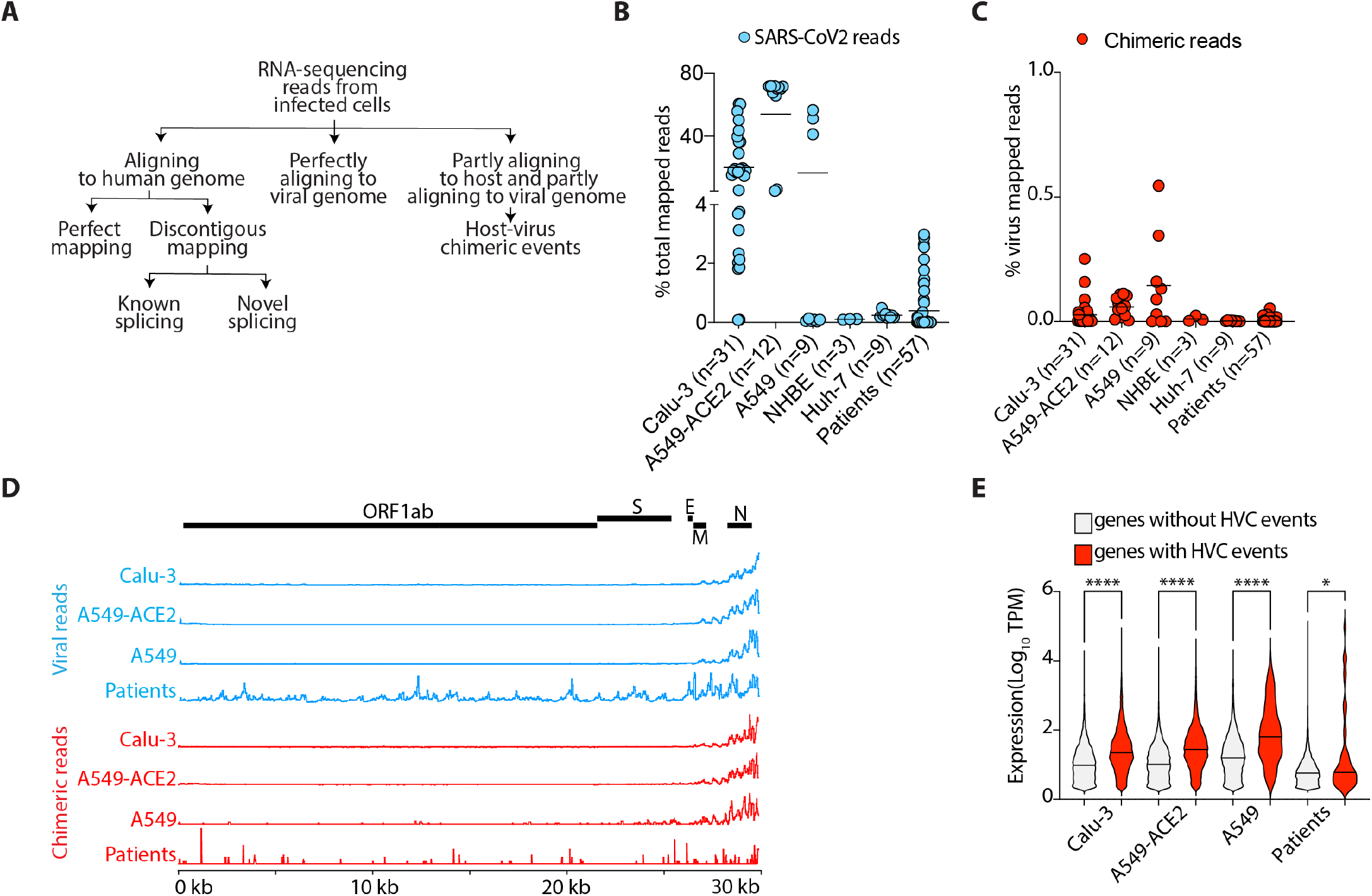
HVC events are detectable in RNA-seq from SARS-CoV2-infected cells but infrequently in samples from COVID-19 patients. (**A**) Schematic presentation of RNA-sequencing data analysis pipeline. (**B**) Viral reads in the indicated SARS-CoV2-infected cells as a proportion of the total reads mapped to the chimeric genome. (**C**) HVC reads in the indicated SARS-CoV2-infected cells as a proportion of the total reads mapped to the SARS-CoV2 genome. (**D**) SARS-CoV2 genome coverage based on reads mapping perfectly to the virus genome (top panel) or to the viral segments of HVC events (bottom panel). (**E**) Violin plots showing the expression of all human genes with or without HVC events in the indicated infected cells. See **Table S1** for the source of Data in this figure. *p<0.05; ****p<0.0001 by Kruskal-Wallis (**E**) and multiple Wilcoxon (**F**) tests and FDR correction.

We next quantified the reads that partly mapped to the human genome and partly mapped to the SARS-CoV2 genome (see **methods**). We found that nearly 0.05-1% of all viral reads are formed of hybrid sequences between host and virus RNAs, a frequency consistent with that recently reported by others ^15,16^ (**Fig. 1C**). Infected A549-ACE2 and Calu-3 cells had the highest percentages of chimeric reads, while others, including normal human bronchial epithelial cells (NHBE) and lung autopsies of patients with COVID-19, had ~1.5-2 orders of magnitude fewer chimeric reads (**Fig. 1C**). Similar percentages of chimeric reads were observed in cells infected with other viruses (**Fig. S1B**).

To test whether there are regions of the viral genome that more frequently participate in chimeric events, we separately aligned the viral reads and the viral fragments of the chimeric reads to the SARS-CoV2 genome (**Fig. 1D**). Consistent with previous studies, we found higher coverage of the 3’ end of SARS-CoV2 genome in sequencing libraries across different cells (**Fig. 1D**, top panel). This portion of the virus encodes the viral N protein. Similarly, we observed that viral fragments from chimeric reads are also biased towards the 3’ end of the SARS-CoV2 genome (**Fig. 1D**, lower panel). This is consistent with a stochastic model in which chimeric events are dependent on the availability of template RNA, i.e., the more viral RNA fragments present the higher the chance of participation in chimeric events. Based on this model, we hypothesized that host fragments participating in chimeric reads will also be over-represented in genes that are more highly expressed. Indeed, we observed that human genes with HVC events are more highly expressed than those without HVC events across all SARS-CoV2 infected cells (**Fig. 1E**). This is exemplified by A549-ACE2 cells, which are transduced to express high levels of angiotensin converting enzyme (ACE) 2, a well-characterized entry receptor for SARS-CoV2. In these cells, ACE2 was one of the top loci participating in chimeric events (**Fig. S1C**).

Collectively, we identified HVC events in RNA-seq from SARS-CoV2 infected cells. We found that these events were very rare in samples from patients with COVID-19 and that there was a correlation between the expression of host and/or viral genes and the frequency of participation in chimera formation. These data indicate that HVC events are either stochastic and biased towards more highly expressed transcripts or authentic biological events involving 3’ regions of the viral genome with specific human genomic elements.

### HVC events are not reproducible and have comparable frequency to artifactual chimeric events

A precise and reproducible junction between host and viral fragments of a chimeric event would be evidence of authentic HVC events occurring as part of the natural life cycle of the virus. To determine whether the junctions of HVC events are precise and reproducible, we compared RNA-seq data from independent studies (**Table S1-2**) and looked for reads that spanned known or novel exon-exon splicing junctions, as well as HVCs (**Figs. 1A-B**). For each cell type, we specifically sourced two or more RNA-seq libraries from independent studies (**Table S1**). As expected, ~90% of known splicing events sourced from RefSeq database were reproducible between independent studies (**Figs. 2A-B**, **S2A**). We also found that nearly one-third of novel (i.e., unannotated) splicing events could also be independently replicated between different studies (**Figs. 2A-B, Fig. S2A**). Conversely, almost none of the exact HVC events were reproducible in independent data sets (**Figs. 2A-B, Fig. S2A**).

**Fig. 2.**
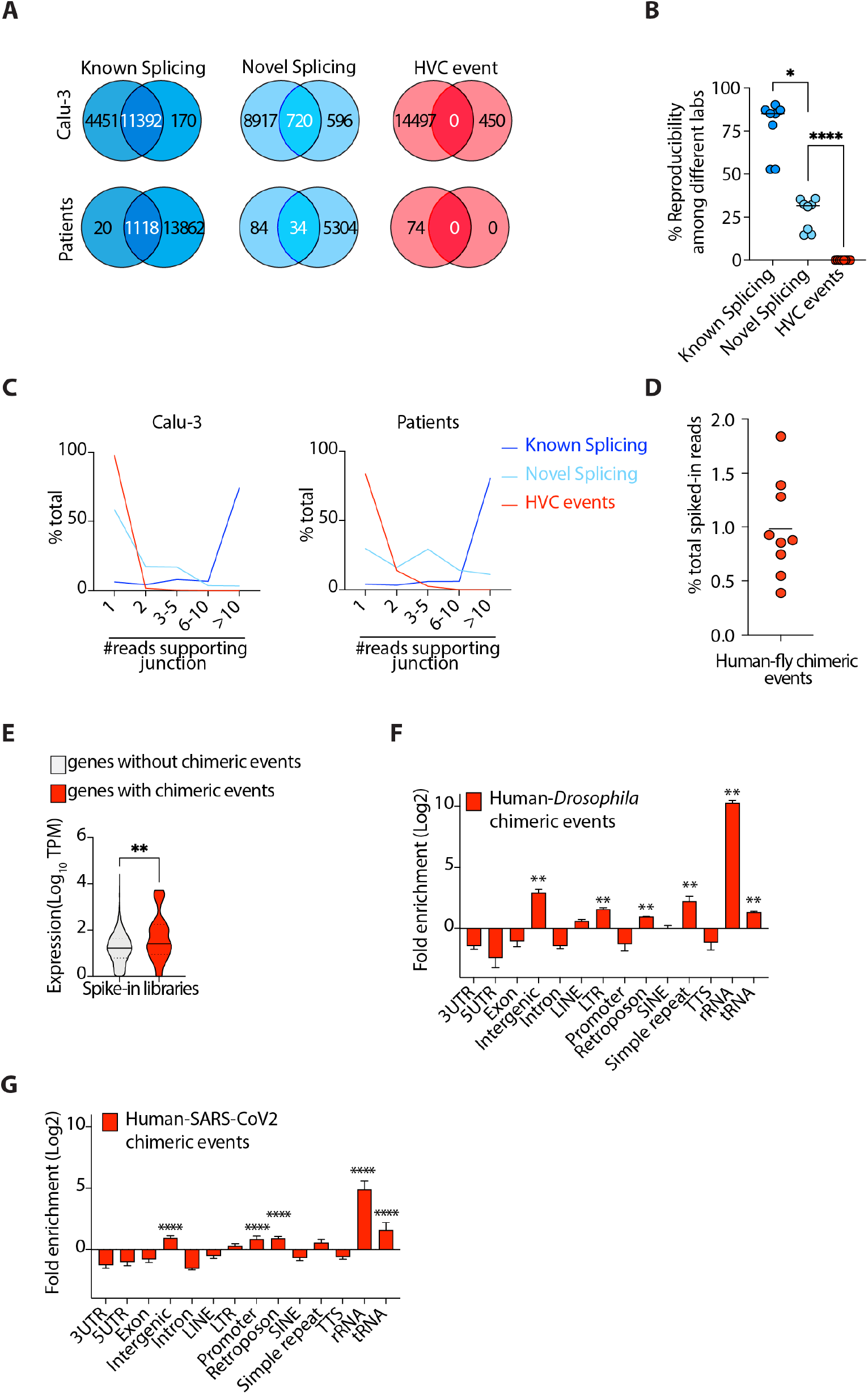
HVC events are not reproducible and have comparable frequencies to artifactual chimeric events. **(A-B)** Representative Venn diagrams (**A**) and cumulative data (**B**) comparing known splicing, novel splicing and HVC events across independent studies (see **Table S1** for the list of independent studies used here). The representative studies in (**A**) are GSE147507 and PRJNA665581 for Calu-3 cells, and GSE147507 and GSE151803 for patient samples. (**C**) Histograms showing the number of reads spanning junctions of the indicated events. (**D**) The fraction of spiked-in *Drosophila* RNA detected to be a chimera with human RNA. Data are from PRJNA311567. (**E**) Violin plots showing expression of all human genes with or without human-*Drosophila* chimeric events. (**F**) Distribution of genomic features in the human segment of human-*Drosophila* chimeric events. (**G**) Distribution of genomic features in the host segment of human-SARS-CoV2 HVC events. *<0.05; **p<0.01; ****p<0.0001 by Wilcoxon test (**B, E**) and FDR correction (**F**).

Another way to determine whether specific HVCs are reproducible is to identify the proportion of unique reads in a given RNA-seq dataset that span the HVC junction. The higher the number of reads the more likely it is that the HVC is not a stochastic event. We thus examined the number of unique reads spanning known splicing, novel splicing and HVC junctions in each RNA-seq dataset (**Fig. 2C**). We found that only 2-15% of HVC events had more than one read spanning their junctions. This is in clear contrast to 90-95% and 40-70% of known and novel splicing events, respectively, that have more than one supporting read (**Fig. 2C, Fig. S2B**).

Our data above indicated that observed HVCs likely represent non-biological artifacts. However, how these artifacts are generated remained unclear. Reverse transcriptase enzymes (RTs) are error-prone and susceptible to a process called random template switching ^18^. In this process RTs synthesizing cDNA infrequently dissociate from their template RNA and associate with a secondary template RNA, resulting in creation of an artifactual fusion cDNA containing both the original template and the secondary RNA. Reverse transcription is one of the main steps in most commonly used RNA-sequencing methods, thus it is conceivable that some of the HVC events are artifacts of RT. To test this, we took advantage of spike-in control libraries that are typically utilized for internal calibration and normalization. In those libraries, a small quantity of RNA from an unrelated species is spiked-in to the RNA of interest, followed by RNA-sequencing library preparation. We sourced existing human RNA-sequencing libraries that harbored spiked-in *Drosophila melanogaster* RNA and were prepared using a common library preparation kit from Illumina. We mapped these libraries to the human-*Drosophila* chimeric genome, using the exact same method that we employed when analyzing the host-virus chimeric genome (see **Methods**). Nearly 5% of all reads were mapped to the *Drosophila* genome. We then identified the fraction of *Drosophila* mapped RNAs that were human-*Drosophila* chimeric. Since there is no actual possibility of biological fusion events between host and spiked-in RNAs, we considered any chimeric reads identified as artifactual. This could therefore determine the expected background level (“noise”) of chimeric events created as artifacts of RT and/or alignment errors. We observed ~1% of all *Drosophila*-mapped reads to participate in chimeric events (**Fig. 2D**). Interestingly, in all libraries analyzed from SARS-CoV2 infected cells, the observed fractions of HVC reads were lower than 1%, indicating that the frequency of HVC events in SARS-CoV2 infected libraries were comparable to the expected background “noise” of chimeric events created as artifacts of RT and/or alignment errors.

We next examined the expression of human genes with and without *Drosophila* chimeras. Similar to what we had observed in SARS-CoV2 infected cells (**Fig. 1E**), human genes with chimeric events were more highly expressed than those without such events (**Fig. 2E**). This was consistent with a stochastic model in which chimeric events are dependent on the availability of template RNA and driven by random RT template switching. Repeat sequences of RNA are known substrates for RT template switching ^18^. To test this, we examined the genomic distribution of the host segments of chimeric events between human and spiked-in *Drosophila* RNA and found that these artifactual chimeric events were, indeed, enriched in RNAs with highly repetitive structures, including rRNAs and tRNAs (**Fig. 2F**). We next sought to determine whether the same observation holds true in virally-infected cells. In RNA-seq libraries of SARS-CoV2-infected cells we found that HVCs were similarly enriched in RNAs with repetitive motifs, including rRNAs and tRNAs, when compared to the total transcriptome (**Fig 2G,** see **methods**).

Collectively, these data indicated that the exact junctions of HVC events were not reproducible between different samples, that the frequency and the genomic distribution of HVC events were comparable to that from artifactual chimeric events generated by RT template switching and that host RNAs partaking in chimera formation were enriched in structures conducive to template switching.

### Experimental enrichment for viral RNA during RNA-seq library construction does not enrich for HVC events

Although viral reads in most infected RNA-sequencing libraries were readily detectable, the fraction of viral reads to total mapped reads was low (**Fig. 1B**), presumably due to heterogenous infectivity rates within cell cultures or patient samples. Thus, it is possible that detection of HVC events and junctional reads is too infrequent to allow robust detection of identical species across different samples. Thus, we developed a technique to experimentally enrich for viral RNAs during RNA-seq library preparation that would also enrich any *bona fide* HVC events as well. To this end, we designed a pool of 30 specific oligonucleotides that spanned the entire SARS-CoV2 genome (**Table S3**). Using these oligonucleotides, we developed a novel methodology to specifically amplify viral RNAs from SARS-CoV2-infected cells and constructed sequencing libraries (schematic in **Fig. 3A**, please also see **methods**). Two types of chimeric events are possible, 5’->3’ host-virus chimeras and 5’->3’ virus-host chimeras. To enrich for viral sequences and ensure “capture” of both types of chimeras, we used two approaches (enrichment methods 1 and 2, respectively, in **Fig. 3A**). To enrich for viral sequences that also contain 5’->3’ host-virus chimeras (enrichment method 1 in **Fig. 3A**), we carried out virus-specific RT to construct cDNA incorporating an Illumina-P5 adaptor sequence and T7 RNA polymerase promoter, followed by second-strand DNA synthesis and *in vitro* RNA transcription using T7 RNA polymerase. RT primed with random hexamer was then carried out to incorporate an Illumina-P7 adaptor sequence, before library amplification by PCR and high throughput Illumina sequencing. To also enrich for viral sequences including 5’->3’ virus-host chimeras (enrichment method 2 in **Fig. 3A**), we performed oligo-dT-primed RT to construct cDNA incorporating an Illumina-P5 adaptor sequence and T7 RNA polymerase promoter, followed by second-strand DNA synthesis and *in vitro* RNA transcription using T7 RNA polymerase. RT primed with virus-specific primers was then carried out to incorporate an Illumina-P7 adaptor sequence, before library amplification by PCR and high throughput Illumina sequencing. Any RNA amplified using this technique would be enriched in viral sequences, including those mapping solely to the viral genome as well as those mapping partially to host as well as viral genomes (HVCs). For comparison, we also prepared cDNAs from RNAs of infected cells without any enrichment (unenriched control, see **methods**).

**Fig. 3.**
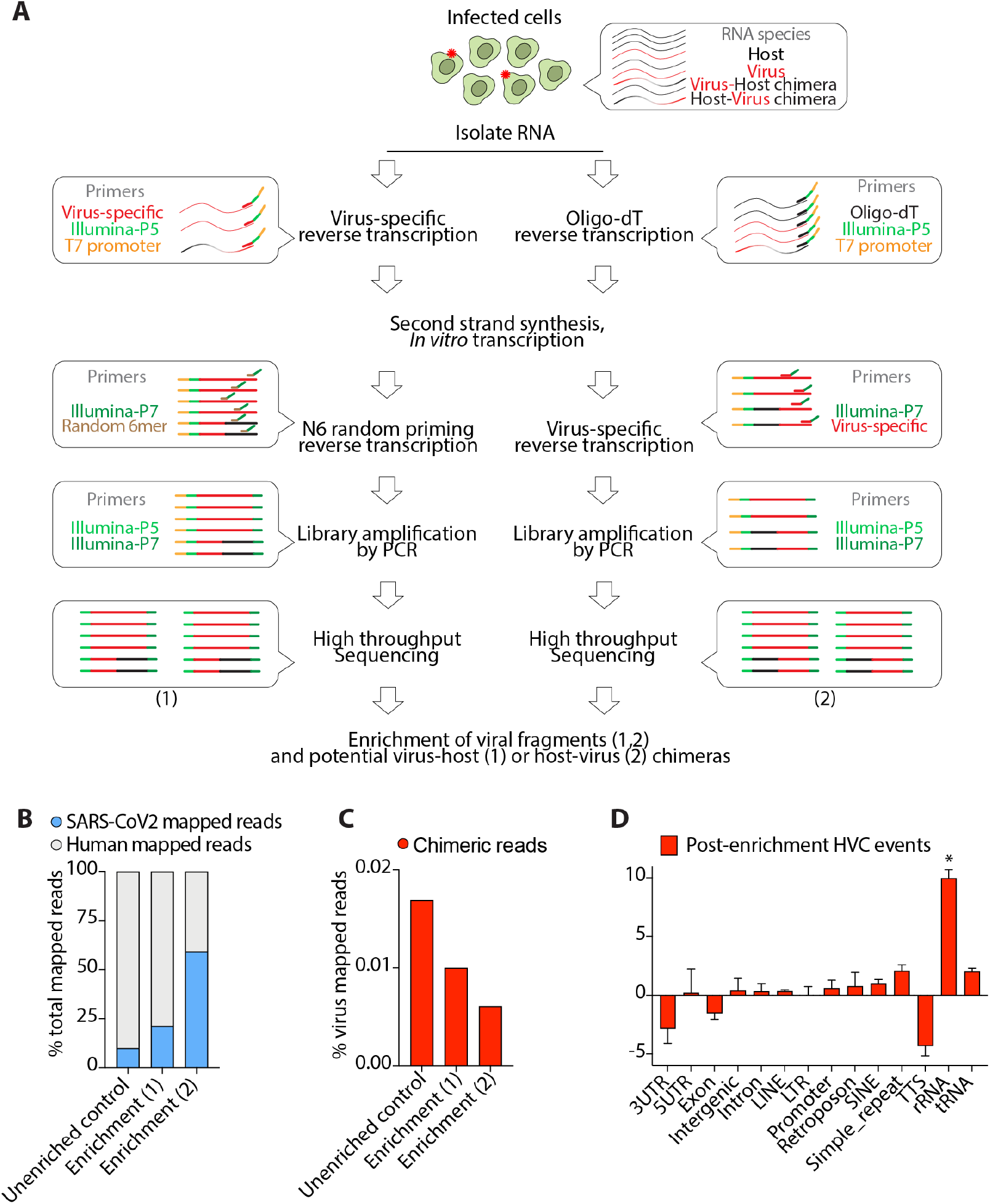
Experimental enrichment for viral containing fragments does not enrich for HVC events. (**A**) Schematic presentation of viral RNA enrichment from infected host cells. Cellular RNA from infected cells comprises host RNA, viral RNA and presumably any fusion RNA between virus and host. A pool of oligo probes that are specific to SARS-CoV2 were used in a series of reverse transcription, *in vitro* transcription and PCR amplification steps to amplify viral RNAs and potential virus-host (1) or host-virus (2) chimeras (see **Methods**). (**B**) Viral reads in the indicated libraries from SARS-CoV2-infected Calu-3 cells as a proportion of the total reads mapped to the chimeric genome. (**C**) HVC reads in the indicated libraries from SARS-CoV2-infected Calu-3 cells as a proportion of the total reads mapped to the SARS-CoV2 genome. (**D**) Distribution of genomic features in the human segment of HVC events detected after enrichment for viral-containing transcripts. * p<0.05 by Wilcoxon test.

To validate this approach, we performed qPCR on sequencing libraries for SARS-CoV2 *N* gene using CDC recommended primer sets. We specifically chose the *N* gene since it is the highest expressed gene and the site of most HVC events. We observed dramatic enrichment (> 30-fold) of viral *N* gene mRNA in enriched libraries compared to the control (**Fig. S3A**). We next performed high throughput sequencing on all libraries and their corresponding controls and performed the same analysis as **Fig. 1**. Consistent with our qPCR data, we found that the total number of reads mapped to the virus genome were much higher in enriched libraries compared to control (**Fig. 3B**), indicating a mean enrichment for viral reads of 2-6-fold. We then compared the total number of HVC events before and after the enrichment. Despite the significant enrichment of viral reads, HVC events were not enriched at all and their frequency remained at <0.05% (**Fig. 3D**), comparable to the expected level from background “noise” denoted previously (**Fig. 2D**). Moreover, the genomic distribution of the host portion of these HVC events (**Fig. 3D**) was similar to those observed from artifactual chimeric events.

Collectively, these data indicate that even after enrichment of transcripts containing viral sequences HVC events remained at the level of noise expected by random RT template switching, which we consider it to be evidence that the observed SARS-CoV2 HVCs are likely artifacts of *in vitro* RNA-seq library construction and unlikely to be *bona fide* events occurring *in vivo*.

## Discussion

Sequencing reads from high throughput assays, if appropriately analyzed, are invaluable resources for identifying novel biological events and provide exceptionally detailed information about host-virus interactions occurring *in vivo*. This is exemplified by the discovery of viral integration as a driver of oncogenesis in HPV-associated cancers ^19^. We have previously used this approach to examine host-virus interactions during EBV infection and to determine how the EBV episome interacts with the human genome ^4,5^. Even in the absence of infection, detailed analyses of RNA-seq data can deliver new insights into cell biology, including, for example, discovery of novel linear or circular genes and isoforms. On the other hand, the technology is extremely sensitive, relies on low fidelity RTs during library preparation and there are many computational challenges during chimeric sequence alignment. Collectively, these can result in the detection of low frequency sequencing reads originating from artifactual events or contaminants (including plasmid vectors). Thus, appropriate positive and negative controls during analysis are essential to distinguish real from artifactual events.

SARS-CoV2 is a good example of a novel pandemic coronavirus of humans that is as yet relatively understudied. In particular, little is known about host-pathogen interactions that determine clinical outcomes, including why some people develop little to no symptoms, while others succumb to life-threatening disease and others progress to chronic disease (“persistent” COVID-19). In this setting, sequencing assays are key methods for uncovering as yet undiscovered mechanisms of pathogenesis. Particular attention has focused recently on reports of HVCs forming between host and viral mRNAs, which have been interpreted as evidence of viral incorporation into the host genome and raised questions about long-term safety of vaccination strategies using viral RNA. Here we found several lines of evidence that indicate that the reported HVC events observed in sequencing libraries are most likely artifactual.

We found that the precise location and the nucleic acid sequence of HVC events are not reproducible across different libraries prepared by different labs. Consistent with previous reports, we concur the viral part of HVC events to be enriched in sequences from the 3’ end of SARS-CoV2 virus. This is the portion of the virus that contains the most highly expressed gene encoding N-protein^20^. Likewise, we also observed that chimeric events incorporated the more highly expressed host genes. A model consistent with these observations is that HVCs are likely the result of stochastic events occurring at the RNA level that incorporate components of more highly expressed transcripts (templates) from both the host and virus.

One of the potential mechanisms that could generate artifactual HVC events is random template switching by RTs used during RNA-seq library preparation to convert RNA to cDNA. RTs are known to occasionally switch from one template to another, thus creating artifactual fusion cDNA. Here we found that ~1% of spike-in control RNAs exhibit chimeric events with host RNA that can only be explained by random template switching during library preparation. This provides an expected level of artifactual chimeric events for the RTs used in common RNA-sequencing library preparation kits (e.g., Superscript II). We found that the frequency of HVC events from all SARS-CoV2 infected samples was below 1%, indicating that these events are likely to be artifacts of RTs. Moreover, repeat sequences of RNA are known substrates for template switching ^18^. Not surprisingly, we found that the host part of HVC events was enriched in RNAs with highly repetitive structures, including rRNAs and tRNAs (**Fig. 1F**). This further supports undesired template switching by RTs as the origin of observed HVC events.

Finally, we developed a novel method to enrich for viral RNA fragments from infected cells. Deploying this method, we found that, although we could enrich for viral transcripts by between >30-fold, the rate of HVCs remained unchanged and at, or below, the expected level of “noise” introduced by *in vitro* RTs. A benefit of our technique is that it is a general method that can easily be used for enrichment of any RNA and its chimeric partners, as long as sequences for oligonucleotide design are known (e.g., a genome build is available). This is particularly useful because cellular RNAs in infected cells typically dominate over RNA derived from infecting pathogens, especially when infection rates and/or viral titers are low. One example for the utility of this method is to help identify “cap-snatching” and “start-snatching” events. In IAV-infected cells, for example, viral transcripts form chimeras with the 5’ portion of host transcripts containing 5’ caps in order to stabilize viral transcripts and create *bona fide* fusion proteins ^21,22^. Although there are computational challenges in aligning sequencing reads if very short fragments (<18 bp) are “snatched” from the host, one would anticipate seeing enrichment of host 5’ UTR elements in HVC events if similar cap-snatching mechanisms were utilized by SARS-CoV2. However, we observed quite the opposite, if any (**Fig. 1F**). In fact, the overall conclusion on successfully enriching for viral RNA reads but observing no enrichment of HVC events above background is that the majority of HVCs are the result of artifacts generated by RT errors during library preparation.

Collectively, our data analyses and experimental findings indicate that currently observed and widely reported HVC events are infrequent, not reproducible, and likely to be an artifact of reverse transcription during RNA-seq library preparation. As anticipated from the cytoplasmic replication stage of positive-strand RNA viruses, viral integration is not expected to be a major pathological factor for SARS-CoV2 and, by extension, not a cause for concern in the use of SARS-CoV2 RNA vaccines.

## Acknowledgments

This research was financed by the National Institute of General Medical Sciences of the NIH (grant R35GM138283 to M.K.) and supported in part by the Intramural Research Program of the NIH, the National Institute of Diabetes and Digestive and Kidney Diseases (NIDDK) (project number ZIA/DK075149 to B.A.), and the National Institute of Allergy and Infectious Diseases (NIAID) (project number ZIA/AI001175 to M.S.L.). D.C. is supported by an NIH Office of Dietary Supplements Research Scholar award.

B.Y., S.C., L.W., B.A. and M.K. analyzed data and wrote the manuscript. C.M., S.C, D.C., D.K. and C.E.W. performed experiments and analyzed data. J.L.T-O., D.C., D.K., M.S.L., M.R.O., C.E.W., B.A. and M.K. provided intellectual input and wrote the manuscript. C.E.W., B.A. and M.K. conceived and supervised the work.

---

**Fig. S1.**
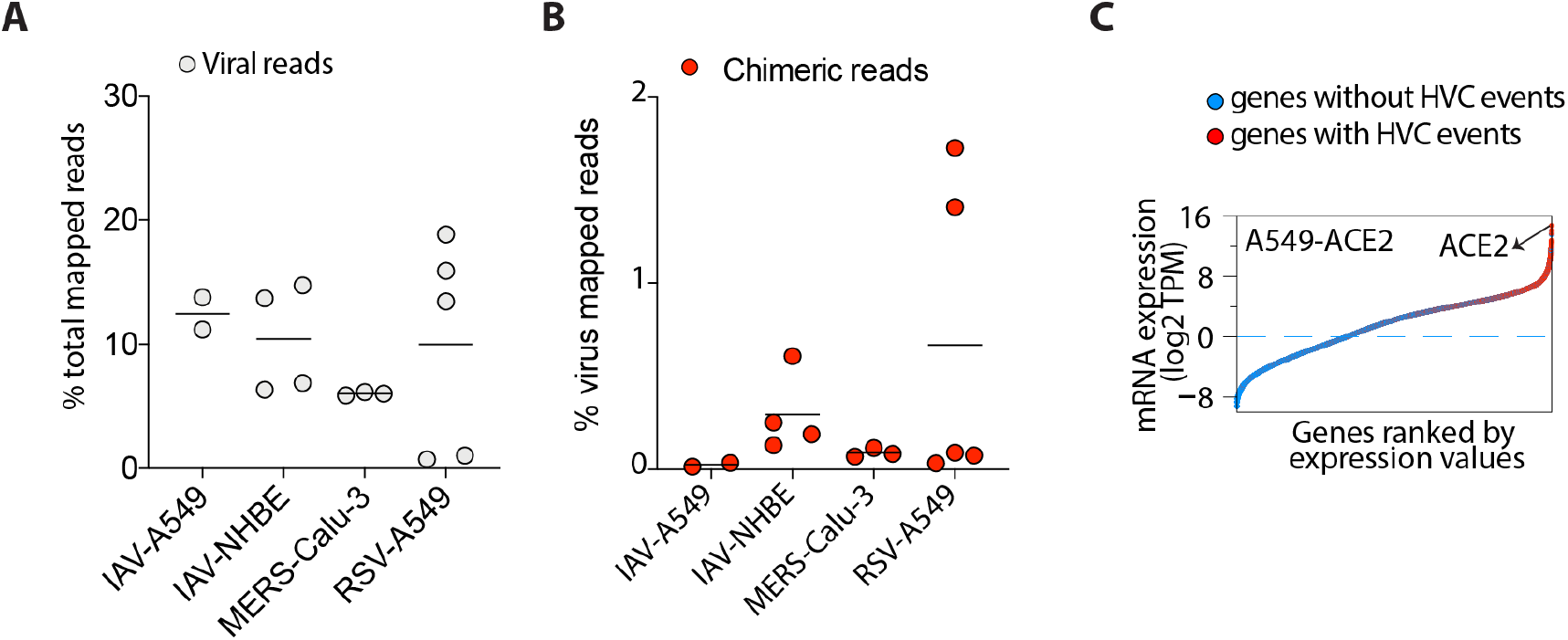
The presence of viral reads and HVC events across infected samples. (**A**) Viral reads in the indicated virus-infected cells as a proportion of the total reads mapped to the chimeric genome. (**B**) HVC reads in the indicated virus-infected cells as a proportion of the total reads mapped to viral genome. (**C**) Dot plots showing the expression of all human genes in SARS-CoV2 infected A549-ACE2 cells ordered by gene expression. Genes with or without HVC events are highlighted with red and blue, respectively.

**Fig. S2.**
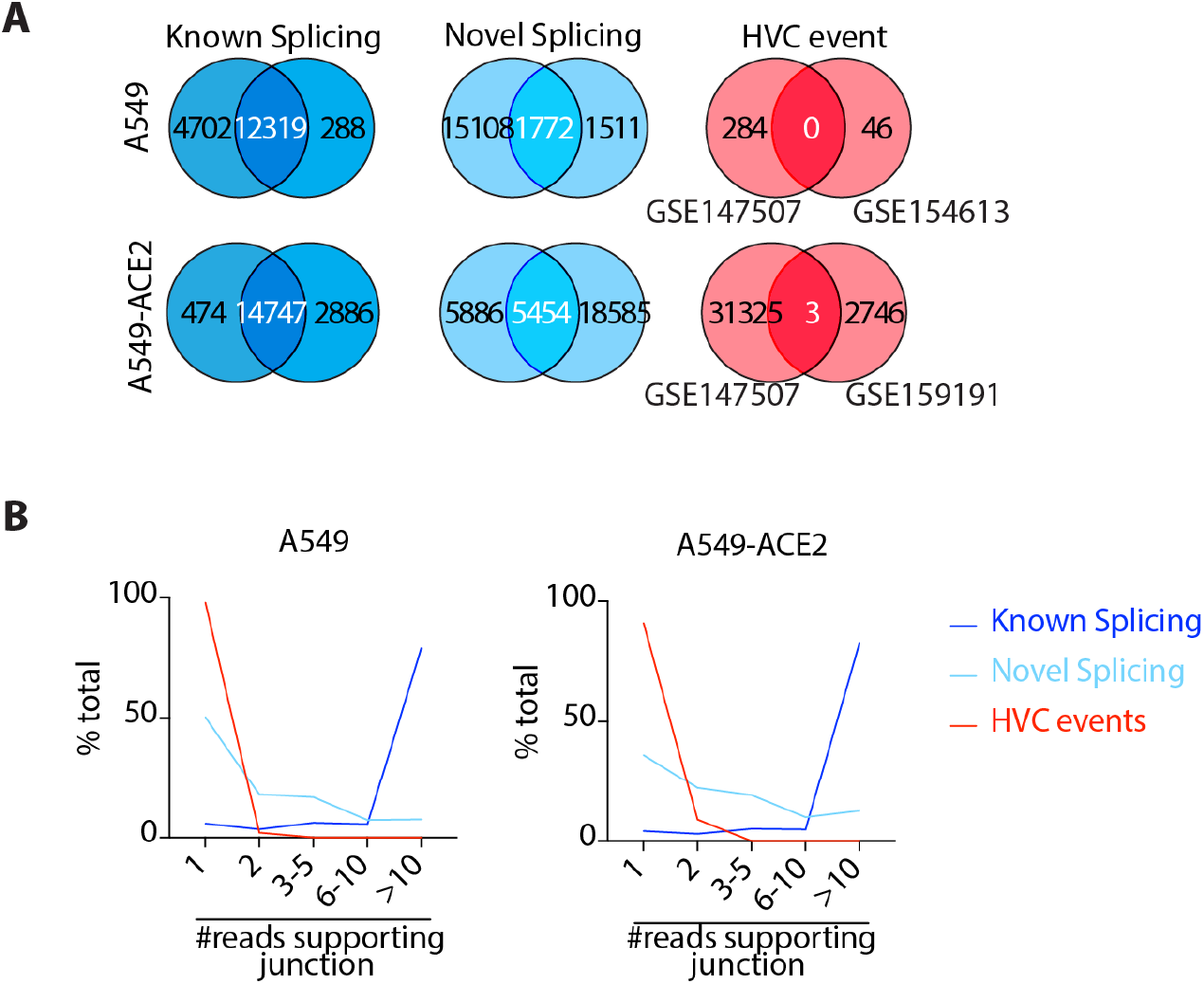
Reproducibility of HVC events in SARS-CoV2 infected A549 and A549-ACE2 cells. (**A**) Venn diagrams comparing known splicing, novel splicing and HVC events in SARS-CoV2 infected A549 (top panel) and A549-ACE2 (bottom panel) cells across independent studies. See **Table S1** for the list of independent studies used here. (**B**) Histograms showing the number of reads spanning the junctions of the indicated events.

**Fig. S3.**
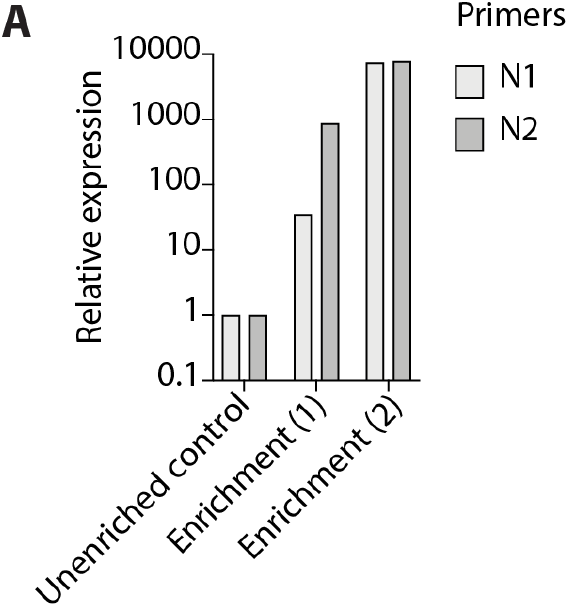
qPCR validation of proposed viral enrichment method. (**A**) Expression of N-protein in control and virus enriched (1 or 2) samples using N1 and N2 qPCR probes recommended by the CDC.

## Tables and Table legends

**Table S1.** SARS-CoV2 infected samples from independent studies used here. See Table S2 for the complete list of individual samples.

| Samples | Accession numbers | Attributions |
| --- | --- | --- |
| A549-ACE2 | GSE147507 | Blanco-Melo *et al.*, *Cell* 2020 |
|  | GSE154613 | Desai *et al.*, *Nat Com* 2020 |
| A549 | GSE159191 | Weingaren-Gabbay *et al.*, *bioRxiv* 2020 |
|  | GSE147507 | Blanco-Melo *et al.*, *Cell* 2020 |
| Calu-3 | GSE147507 | Blanco-Melo *et al.*, *Cell* 2020 |
|  | PRJNA665581 | Banerjee *et al.*, *Cell* 2020 |
|  | GSE148729 | Wyler *et al.*, *bioRxiv* 2020 |
| COVID-19 Patients | GSE147507 | Blanco-Melo *et al.*, *Cell* 2020 |
|  | GSE151803 | Schwartz *et al.* |
|  | GSE150316 | Desai *et al.*, *Nat Com* 2020 |

**Table S2. The detailed information of all RNA-seq libraries used in this study. (A-B)** The detailed information of RNA-seq libraries from SARS-CoV2 infected (A) or other virally infected cells (B) used in this study. The library size, the total number of reads mapped to the human-virus chimeric genome genome, viral genome or HVC reads are reported.

**Table S3. Primers/oligoes used in this study. (A)** Primers used for qPCR validation. (**B-C**) Oligos used in viral fragment enrichment method 1 (B) and 2 (C), respectively.

## Methods

### Cell culture and viral infections

Human adenocarcinomic lung epithelial (Calu-3) cells (ATCC, HTB-55) were cultured in Dulbecco’s Modified Eagle Medium (DMEM, GIBCO) supplemented with 10% Fetal Bovine Serum (FBS, Corning), HEPES, non-essential amino-acids, L-glutamine and 1X Antibiotic-Antimycotic solution (Gibco). All cells were maintained at 37°C and 5% CO2. SARS coronavirus 2 (SARS-CoV-2), isolate USA-WA1/2020 (NR-52281) was obtained from Bei resources and was propagated in Vero E6 cells in DMEM supplemented with 2% FBS, 4.5 g/L D-glucose, 4 mM L-glutamine, 10 mM Non-Essential Amino Acids, 1 mM Sodium Pyruvate and 10 mM HEPES. Infectious titers of SARS-CoV-2 were determined using TCID50 method (Reed and Muench). Fifty thousand Calu-3 were seeded in 48 well plate and allowed to form 80% confluent monolayer. SARS-CoV-2 virus was pre-treated with porcine trypsin (10ug/ml) for 15 minutes at 37 degrees. Cells were then infected with the pre-treated virus prep at a multiplicity of infection (MOI) of 10 for 1h in culture media (final concentration of trypsin on cells was 2ug/ml). After absorption, virus inoculum was removed and replaced with fresh culture media. 48hrs post infection, cells were harvested and RNA was isolated. Briefly, infected cells were lysed in TRIzol (Invitrogen) and RNA was extracted using Direct-zol RNA Miniprep kit (Zymo Research) as per manufacturer’s instructions. Experiments using SARS-CoV-2 were performed at the University of Michigan under Biosafety Level 3 (BSL3) protocols in compliance with containment procedures in laboratories approved for use by the University of Michigan Institutional Biosafety Committee (IBC) and Environment, Health & Safety (EHS).

### Library preparation and Virus Enrichment Assay

To experimentally enrich for viral RNAs from total RNA of SARS-CoV2 infected cells prior to RNA-seq library preparation, we developed a series of *in vitro* amplification steps using SARS-CoV2 virus specific primers (VSP) as following. The VSP pool contained ~30 oligonucleotides that span all SARS-CoV2 genes (at least one oligo per gene and nearly 1 oligo per 1kb of the genome). Given our goal to additionally enrich for potential HVC events, we used two approaches, enrichment methods 1 (5’->3’ host-virus chimeras) and enrichment method 2 (5’->3’ virus-host chimeras) (**Fig. 3A**).

#### First strand cDNA synthesis reaction

To capture and enrich for viral, virus-host or host-virus transcripts, we first set up a 20 μl reverse transcription (RT) reaction using 100 ng of mRNA isolated from SARS-CoV2 infected using Superscript III reverse transcriptase. We used 2 pmol T7-P5-VSP oligo pool (for enrichment 1) or 50 pmol of T7-P5-Oligo dT (for enrichment 2) as “gene-specific primer” for RT reaction (see **Table S3**). We also incorporated a T7 promoter and Illumina P5 sequence at the 5’ end of every oligo as shown in the schematic **Fig. 3A**. After combining all the components as per the recommended protocol (Thermo Fisher, Catalog# 18080044 and 18064014), we incubated the entire reaction mixture at 25°C for 15 minutes followed by 50°C for 30 minutes for SSIII. Next, we inactivated the reaction by heating at 70°C for 15 minutes. To remove RNA/complementary RNA to the cDNA, we added 1 μl of PureLink™ RNase A (20 mg/mL) (Invitrogen Catalog# 12091021) and 1 μl (5 units) of RNase H (abmgood catalog# E018) and incubated at 37°C for 1 hour. We then purified cDNA using 1X Mag-Bind TotalPure NGS (Omega Bio-tek, Catalog# M1378-01) as per manufacturer’s instructions and eluted the cDNA in 15 μl of sterile water. To further remove the excess single-stranded short oligos, we treated the purified RT reaction with 1 μl of Exonuclease I (NEB, Catalog# M0293L) at 37°C for 30 minutes. Next, we added excess sterile water to sample to get a total volume of 40 μl. The reaction was them purified with 1X Mag-Bind TotalPure NGS beads and eluted in 15 ul of sterile water.

#### Second strand cDNA synthesis and In vitro Transcription

Following this, we performed second strand synthesis using the NEBNext Ultra II Non-Directional RNA second strand synthesis module as per the suggested protocol (NEB, Catalog# E6111L). The synthesized DNA was purified via 1X Mag-Bind TotalPure NGS beads and eluted in ~12 ul of sterile water. 10 ul of this was then used as an input for T7 polymerase mediated In vitro Transcription (IVT) using the NEB HiScribe T7 High Yield RNA Synthesis Kit (NEB, # E2040S). Briefly, all the components were mixed as mentioned in the kit protocol and incubated at 37°C (lid at 50°C) for 16 hours. The reaction was eluted in 20 μl of sterile water after a round of 1X Mag-Bind TotalPure NGS bead cleanup. This newly transcribed RNA was quantified using a Nanodrop and to improve the hybridization kinetics and enhance signal, 500ng of the amplified RNA was fragmented by RNA Fragmentation reagent in a total reaction volume of 10 μl as per specifications (Thermo Fisher, Catalog# AM8740).

#### Final reverse transcription and PCR enrichment of the library

Next, to generate final enriched libraries, we performed reverse transcription of the fragmented RNA with 50 pmol of P7-N6 for enrichment (1) and 2 pmol of the P7-VSP primer pool for enrichment (2,) respectively (see **Table S3**) using SSIII reverse transcriptase as per the steps mentioned above. After RT, the reaction mixture was purified using 1X Mag-Bind TotalPure NGS beads and eluted in 20 μl of sterile water. 5 μl of this RT reaction was saved for running on a Bioanalyzer and to perform a qRT-PCR validation assay. The remaining 15 μl was used to PCR amplify the library by using high-fidelity Q5 DNA Polymerase (NEB, Catalog# M0491L) for 16 cycles using Universal primer and unique indices (NEB Catalog# E7335L, E7500L) in a total of in a 50 μl reaction volume. Finally, the amplified and enriched library was purified using the 0.8X Mag-Bind TotalPure NGS beads, quantified by Bioanalyzer/Tape station and then sequenced using Illumina platform.

### qRT-PCR validation assay

The enrichment of viral genes was determined by performing a qRT-PCR assay on the libraries generated. Briefly, the cDNA generated by RT, prior to library amplification by Q5-PCR was diluted 10 −20 fold and used to amplify target gene ‘N’ of SARS-Cov2 using CDC recommended primers for 2019-nCoV_N1-F: 5’-GAC CCC AAA ATC AGC GAA AT-3’, 2019-nCoV_N1-R: 5’ TCT GGT TAC TGC CAG TTG AAT CTG-3’, 2019-nCoV_N2-F: 5’ TTA CAA ACA TTG GCC GCA AA-3’, 2019-nCoV_N2-R: 5’ GCG CGA CAT TCC GAA GAA3’ and UBC as the housekeeping gene, UBC-F: 5’-CCT GGA GGA GAA GAG GAA AGA GA-3’ and UBC-R: 5’-TTG AGG ACC TCT GTG TAT TTG TCA A-3’. The qRT-PCR was performed on Bio-Rad CFX connect. All experiments were performed in independent triplicates in total reaction volumes of 15 μl using PowerUp SYBR Green Master Mix (Applied Biosystems, Catalog# A25778). The expression was calculated by the 2^-dCt^ method and normalized to that of indicated housekeeping gene in the same sample.

### Host-virus chimeric read analysis

The raw sequencing files were download from the Sequence Read Archive (SRA) as shown in the **Table S1**. Fastqc (v0.11.7) was used for data quality control. Sequencing reads were aligned as single end to the chimeric genome of human (hg38) and SARS-CoV2 (NC_045512.2) using STAR aligner (v2.7.7a). For the analyses of the other viruses, the influenza A virus (IAV) genome (A/Puerto Rico/8/1934 (H1N1), GCA_000865725.1), middle east respiratory syndrome (MERS) genome (NC_019843.3) and the Respiratory syncytial virus (RSV) genome (A2 strain, M11486) were all sourced from NCBI.

To estimate the background level of chimeric reads in RNA-seq libraries, a fruit-fly RNA spikein control libraries (PRJNA311567) were used. Briefly, a chimeric genome between human (hg38) and fruit-fly chr4 (dm6) was constructed and the sequencing reads were aligned by the STAR aligner using the same parameters as --outFilterMultimapNmax 1 --outFilterMismatchNmax 3 --chimSegmentMin 30 --chimOutType Junctions SeparateSAMold WithinBAM SoftClip --chimJunctionOverhangMin 30 --chimScoreMin 1 --chimScoreDropMax 30 --chimScoreJunctionNonGTAG 0 --chimScoreSeparation 1 --alignSJstitchMismatchNmax −1 −1 −1 −1 --chimSegmentReadGapMax 3.

The known annotated and novel unannotated splicing junctions were extracted from the STAR output as positive controls. The chimeric junctions for human-virus and human-fly were extracted from the STAR chimeric output. The unique chimeric junctions were considered as our chimeric events. To estimate the reproducibility, for each independent study and each cell type, the number of unique junctions were extracted. For every cell type, the mean value of overlapping junctions from two independent studies to the number of junctions in each study was recorded as the reproducibility.

To examine the genomic features of the HVC reads, HOMER (v4.11) annotatePeaks.pl was used to annotate the HVC junctions and the corresponding RNA-seq library. In brief, reads in each RNA-seq library were converted to genomic regions by bamTobed (bedtools, v2.30.0) and the unique regions were kept using the following command”sort -k1,1 -k2,2n | uniq”. The reported “Log2 Ratio (obs/exp)” for each annotation (e.g tRNA, LTR) were compared between HVC junctions and the corresponding RNA-seq library. Mann-Whitney’s U test was used for statistical analysis.

